# A chemosensory circuit for microbial metabolite-guided attraction in *C. elegans*

**DOI:** 10.64898/2026.11.18.767703

**Authors:** Tsung-Lin Tsai, Rocio Renee Rubiano, Alejandro Aballay

## Abstract

Animals inhabit rich microbial landscapes in which volatile cues signal both nutrient opportunities and infection risk. In the nematode *Caenorhabditis elegans*, chemosensation guides foraging decisions among diverse bacteria, yet the mechanisms underlying naive attraction to pathogens remain unclear. Here we demonstrate that *C. elegans* exhibits a strong olfactory attraction to *Salmonella enterica* and closely related Enterobacteriaceae isolates. This attraction relies on the bacterial production of isoamyl alcohol (IAA) and hydrogen sulfide (H₂S), two volatiles that signal the presence of metabolically active Enterobacteriaceae, causing the animal to approach *S. enterica* despite its pathogenicity. Sensory neuron classes ASE, ASI, and AWC mediate the response, and the behavioral preference is sharpened by prior experience. Animals fed *S. enterica* mutants defective in leucine biosynthesis (IAA production) or sulfur metabolism (H₂S production) show greatly reduced attraction, while exogenous IAA or H₂S rescues the effect. These results reveal a defined chemosensory circuit through which conserved microbial metabolites modulate host behavior, uncovering a form of behavioral adaptation that integrates microbial cues to optimize foraging under pathogenic uncertainty.

**SIGNIFICANCE STATEMENT:** Animals locate food by smell. Among the odors they detect are the volatile chemicals released by bacteria as they grow. These same chemicals are also produced by harmful bacteria, so an odor that signals a meal can also lead to infection. We show that *C. elegans* is strongly attracted to the human pathogen *Salmonella enterica*, and that this attraction is driven by two ordinary bacterial metabolites, isoamyl alcohol and hydrogen sulfide, the gas responsible for the smell of rotten eggs. A defined set of sensory neurons reads these cues, and prior infection can change their meaning. Bacterial pathogens can therefore exploit a conserved attraction to the chemistry of microbial growth, turning a foraging signal into a route to infection.

## INTRODUCTION

From humans avoiding the smell of spoiled food to a nematode navigating a decomposing substrate, animals rely on the volatile chemicals released by microbes to decide what is safe to eat. This strategy is powerful but imperfect. The bacterial metabolism that generates the appealing odors of a nutrient-rich food source also generates the repellent odors of decay, and pathogens can emit the very same cues as harmless microbes. For the nematode *Caenorhabditis elegans*, which naturally inhabits decomposing plant material and other colonized substrates, microbial community composition changes rapidly, and the animals must track these changes to survive (1–3). In this context, olfactory cues are central to bacterial discrimination and habitat selection, because they provide long-range information about microbial identity and metabolic state before direct contact occurs (4, 5). Many of these cues originate from benign or nutritional bacterial associates, yet pathogens can emit similar chemical signals. The nematode must therefore distinguish benefit from risk in real time based on overlapping sensory inputs.

The natural microbiota of *C. elegans* includes diverse Proteobacteria, Bacteroidetes, Firmicutes, and Actinobacteria, with Enterobacteriaceae featuring prominently among isolates from rotting fruits and vegetation (6–8). These bacterial communities are recruited from the surrounding substrate, yet their composition in the gut differs from that of the environment, indicating selective colonization and persistence (2, 7). *Salmonella enterica* and related Enterobacteriaceae can colonize the *C. elegans* intestine, proliferate, and alter host physiology and survival, establishing the nematode as a tractable model to study *S. enterica* pathogenesis and host responses (9–12). Previous microbiome and pathogenesis studies have shown that gut-associated bacteria differ in metabolic capacity, colonization dynamics, and impact on host life-history traits, suggesting that *C. elegans* responds selectively to microbial metabolites that predict resource quality and infection risk (6–8). However, the olfactory and neuronal basis of *C. elegans* responses to *S. enterica* and other Enterobacteriaceae, and how these responses are shaped by experience, remain poorly understood.

In many host-microbe systems, bacterial volatile metabolites function as ecological signals that report microbial density, growth, and nutrient status. Foraging animals use such compounds to locate nutrient-rich patches or detect abundant microbial biomass, often by sampling conserved metabolic products that are shared across taxa (6, 8). In *C. elegans*, the AWA and AWC olfactory neurons detect a wide panel of attractive odors, including short-chain alcohols and ketones produced during bacterial fermentation (4, 5). Two metabolites of particular interest, isoamyl alcohol (IAA) and hydrogen sulfide (H₂S), arise from branched-chain amino acid metabolism and sulfur reduction, respectively, and are enriched in bacterial lineages that thrive in decomposing substrates, including many Enterobacteriaceae (4, 5, 13, 14). IAA is a well-characterized AWC ligand that elicits robust attraction and undergoes experience-dependent modulation, whereas H₂S is a gaseous metabolite produced by multiple bacterial pathways and has emerged as an important signal in microbial physiology and host interactions (4, 5, 13, 14). Their production by Enterobacteriaceae suggests that they may act as reliable cues of microbial activity and metabolic vigor. An animal that orients toward these volatiles can therefore locate dense bacterial substrates, even at the cost of encountering potentially pathogenic strains.

Here we show that attraction of *C. elegans* to *S. enterica* depends on ASE, ASI, and AWC sensory neurons and that the valuation of microbial odors is strongly modulated by prior experience. We identify IAA and H₂S as two conserved volatiles that mediate this behavior. By combining bacterial genetics, defined microbial isolates, and neuronal manipulations, we define a pathway in which conserved microbial metabolites act through specific chemosensory circuits to shape host choice behavior. These findings show that *C. elegans* integrates olfactory cues, learning, and ecological context to navigate complex microbial environments.

## RESULTS

### *Salmonella enterica* odor elicits *C. elegans* attraction that is experience-dependent

Although pathogenic bacteria often elicit avoidance in *C. elegans*, behavioral responses to pathogens are not uniform. In some cases, avoidance emerges only after a prolonged pathogen exposure, when intestinal colonization and distension generate internal danger signals that modify behavior (15–18). However, despite its ability to colonize the intestine and induce distension, the pathogen *S. enterica* does not elicit avoidance (12, 19). This observation raises the possibility that *S. enterica* emits volatile cues that attract the animals, overriding avoidance responses normally associated with intestinal infection.

To test whether attraction to *S. enterica* can be driven solely by volatile cues, we used a spatial two-choice olfactory chemotaxis assay in which animals choose between two otherwise identical *Escherichia coli* lawns differing only in airborne exposure. Preference was quantified using a choice index (CI). Gravid adult wild-type (N2) animals exhibited a significant preference for the *E. coli* lawn paired with *S. enterica* odor (Figure 1A), demonstrating that volatile compounds alone are sufficient to drive attraction. These results show that *S. enterica* produces volatile cues that attract *C. elegans*.

**Figure 1.**
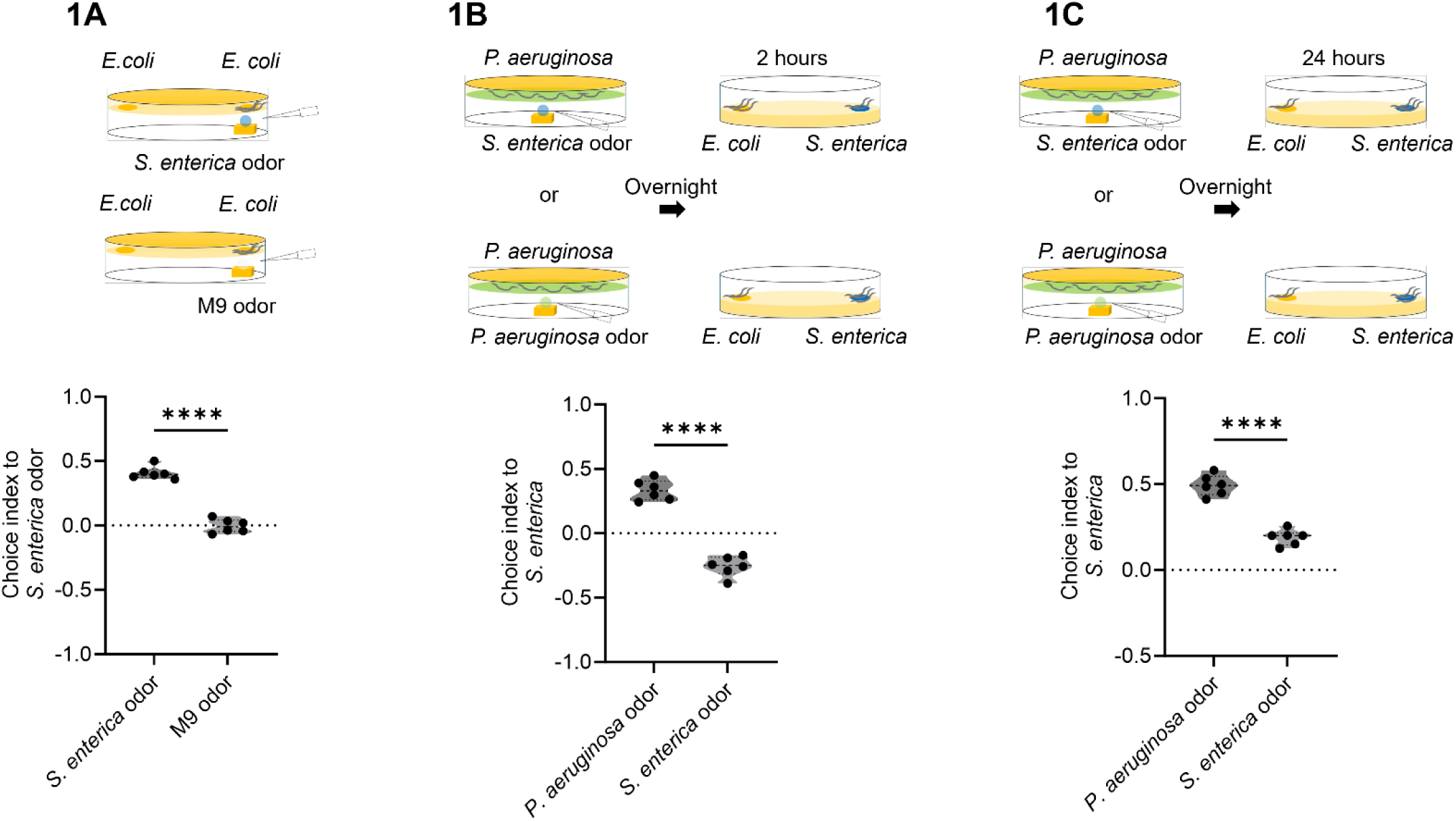
Odor exposure influences *C. elegans* preference for *S. enterica*. The Choice Index (CI) was calculated as (number of worms on *S. enterica* odor side – number on control side) / (number of worms on *S. enterica* odor side + number on control side). CI values above zero indicate preference for *S. enterica* odor. (A) Gravid adult wild-type (N2) *C. elegans* were tested in a two-choice assay between a control *E. coli* lawn and one supplemented with either *S. enterica* odor. *t*-test was used and P values for CI of *S. enterica* odor were calculated with respect to the M9 odor control. ****P < 0.0001, ***P < 0.001, **P < 0.01, and *P < 0.05. ns, non-significant. (B) Gravid adult wild-type (N2) *C. elegans* were fed *P. aeruginosa* and then exposed overnight to either *S. enterica* odor or *P. aeruginosa* odor. The next day, worms were tested in a two-choice assay between *E. coli* and *S. enterica* lawns at 2 hours. (C) Gravid adult wild-type (N2) *C. elegans* were fed *P. aeruginosa* and exposed overnight to either *S. enterica* odor or *P. aeruginosa* odor. The next day, worms were tested in a two-choice assay between *E. coli* and *S. enterica* lawns at 24 hours. Preference was quantified using a Choice Index (CI), where values greater than zero indicate preference for *S. enterica* and values less than zero indicate preference for *E. coli*. *t*-test was used and P values for CI of *P. aeruginosa* odor were calculated with respect to the *S. enterica* odor. ****P < 0.0001, ***P < 0.001, **P < 0.01, and *P < 0.05. ns, non-significant.

Our finding that *S. enterica* odor elicits attraction indicates that this pathogen emits airborne metabolites that drive attraction behavior in *C. elegans*. In natural microbial landscapes, bacterial volatile organic compounds function as long-range foraging cues that guide nematodes toward metabolically active food sources (3, 20). However, many odor-emitting bacteria are also pathogenic. Thus, attraction to microbial volatiles presents an ecological trade-off, as cues that predict nutrient availability may simultaneously signal infection risk. In *C. elegans*, avoidance of bacterial pathogens often emerges only after prolonged exposure, indicating that behavioral responses to pathogens are not determined solely by innate olfactory input but can be reshaped by experience (21).

Determining whether attraction to *S. enterica* odor is fixed or experience-dependent is important for understanding how microbial cues guide foraging decisions. To test this, gravid adults were conditioned for 24 h on *Pseudomonas aeruginosa* PA14, an opportunistic pathogen that induces learned avoidance following intestinal colonization (21, 22). During conditioning, animals were infected with *P. aeruginosa* and simultaneously exposed overnight to volatiles from either *P. aeruginosa* or *S. enterica*. Animals were then transferred to a two-choice assay between *S. enterica* and *E. coli* lawns. *P. aeruginosa*-infected animals exposed to *P. aeruginosa* odor exhibited attraction to *S. enterica*, whereas *P. aeruginosa*-infected animals exposed to *S. enterica* odor displayed robust avoidance (Figure 1B). The difference in odor preference between the two conditioning groups emerged by 2 h (Figure 1B) and persisted for at least 24 h (Figure 1C), indicating durable revaluation of odor preference. At 24 h, animals conditioned with *S. enterica* odor showed reduced, rather than fully reversed, attraction (Figure 1C).

Because all animals experienced *P. aeruginosa* infection, the different behavioral outcomes reflect associative learning. Only animals in which *S. enterica* volatiles were paired with *P. aeruginosa* infection subsequently avoided *S. enterica*, demonstrating that odor preference is reassigned through experience. Thus, attraction to *S. enterica* odor is not fixed but dynamically updated, supporting a model in which bacterial volatile metabolites function as ecological signals whose behavioral meaning is reshaped by prior microbial encounters.

### An ASE–ASI–AWC chemosensory circuit mediates experience-dependent revaluation of *S. enterica* odor in *C. elegans*

The experiments above establish that *S. enterica* emits an attractive odor that can be dynamically reassigned when animals are exposed to the odor in the presence of *P. aeruginosa*. The neuronal mechanisms underlying this change in odor preference remain unknown. Experience-dependent plasticity requires detecting the odor and integrating that signal with changes in host physiology induced by *P. aeruginosa* exposure. Whether this reassignment occurs within canonical volatile-sensing neurons or emerges from distributed chemosensory integration is unclear.

In *C. elegans*, amphid sensory neurons are functionally specialized. For example, ASE mediates chemotaxis to soluble cues (23), ASI regulates neuroendocrine and pathogen-responsive states (22), and AWB and AWC detect volatile odorants (20). Thus, reassignment of *S. enterica* odor preference could reflect plasticity within volatile-sensing neurons (AWB/AWC), modulation by infection-responsive neurons (ASE/ASI), or coordinated integration across these pathways.

To define the neuronal substrate underlying this revaluation, we investigated whether the ASE, ASI, AWB, and AWC sensory neurons are required for experience-dependent reassignment of *S. enterica* odor preference. Wild-type animals and strains defective in ASI, ASE, AWB, or AWC function were conditioned overnight on *P. aeruginosa* in the presence of either *S. enterica* odor or *P. aeruginosa* odor and subsequently tested in two-choice assays. ASE(-), ASI(-), and AWC(-) animals displayed significantly reduced aversion to *S. enterica* relative to *E.coli* (Figure 2A), in contrast, AWB ablation had no significant effect (Figure 2A). We next examined behavioral responses toward *P. aeruginosa* versus *E. coli* following exposure to *P. aeruginosa* lawns and *S. enterica* odors. ASI(-), AWB(-), and AWC(-) animals displayed significantly reduced aversion to *P. aeruginosa* relative to wild-type animals (Figure 2B), consistent with previous reports (24, 25).

**Figure 2.**
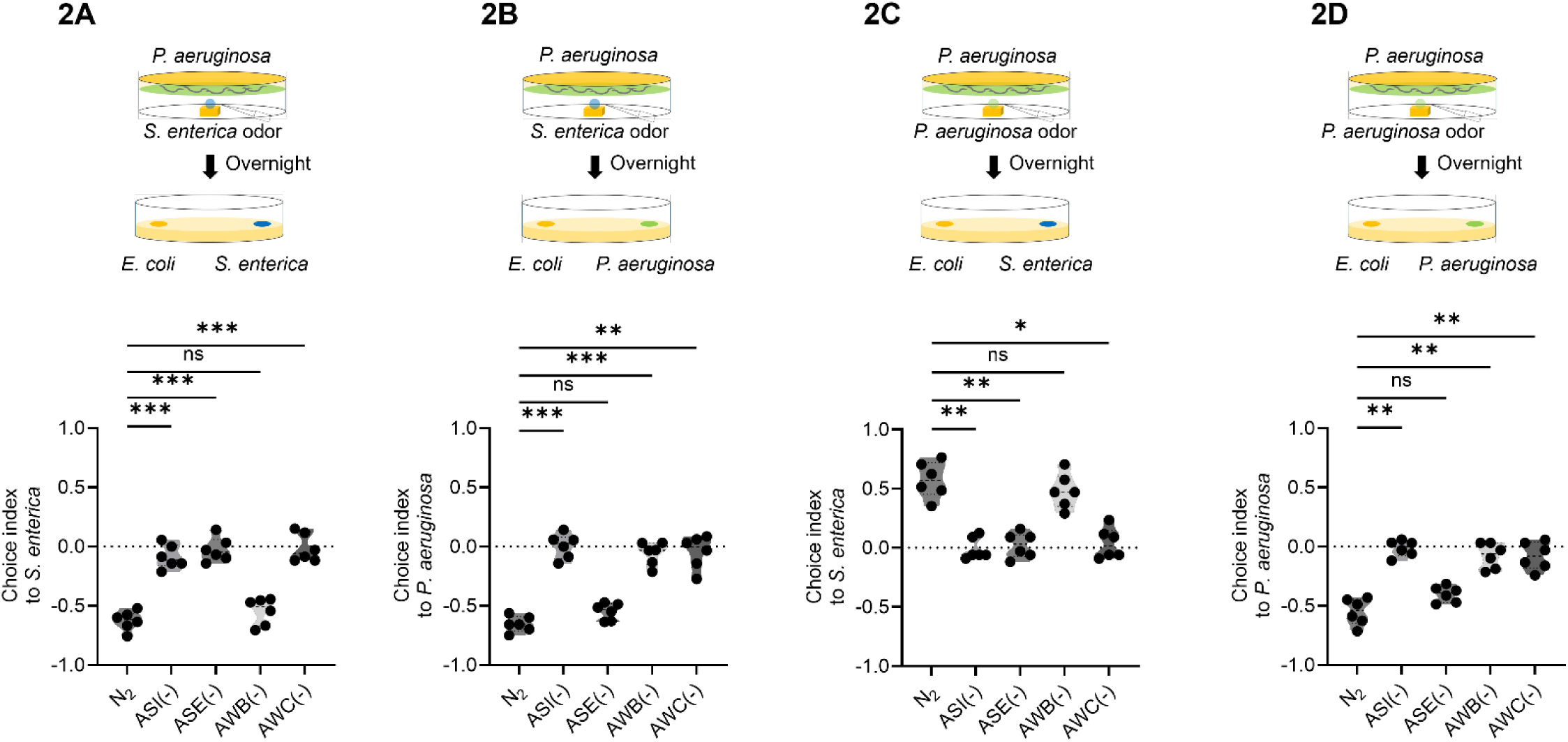
Gravid adult wild-type (N2) and sensory neuron–ablated animals [ASI (-), ASE (-), AWB (-), AWC (-)] were fed overnight with either *P. aeruginosa* along with *S. enterica* odor (2A and 2B) or *P. aeruginosa* along with *P. aeruginosa* odor (2C and 2D). The following day, animals were tested in a two-choice assay between *S. enterica* and *E. coli* lawn (2A and 2C) or *P. aeruginosa* and *E. coli* lawn (2B and 2D). A positive choice index indicates attraction, while a negative index indicates aversion. One-way ANOVA with Tukey’s multiple comparisons test was used and P values for neuron–ablated animals were calculated with respect to the N_2_ control. ****P < 0.0001, ***P < 0.001, **P < 0.01, and *P < 0.05. ns, non-significant.

Finally, we examined whether neuronal ablation affected *P. aeruginosa* avoidance following simultaneous *P. aeruginosa* odor and lawn conditioning. ASE(-), ASI(-), and AWC(-) animals displayed significantly reduced attraction to *S. enterica* (Figure 2C), whereas ASI(-), AWB(-), and AWC(-) animals displayed significantly reduced aversion to *P. aeruginosa* (Figure 2D). While the requirement for ASI in *P. aeruginosa* avoidance is consistent with previous reports (24, 25), its requirement for attraction to *S. enterica* reveals a previously unreported role for ASI in mediating attraction. Together, these data identify an ASE–ASI–AWC chemosensory circuit required for experience-dependent revaluation of *S. enterica* odor in *C. elegans*.

### Conserved Enterobacteriaceae Volatiles Drive Olfactory Attraction in *Caenorhabditis elegans*

The sustained attraction of *C. elegans* to *S. enterica* is surprising because this bacterium is pathogenic and can colonize and distend the intestine. Moreover, as shown above, the behavioral response to *S. enterica* odor can be reassigned when that odor is encountered in an aversive context associated with *P. aeruginosa*. This raises an important question about the nature of the signal driving attraction. One possibility is that animals recognize volatile cues specific to *S. enterica*. Alternatively, attraction may arise because the pathogen produces conserved metabolic volatiles that normally signal the presence of metabolically active bacterial food sources. In natural environments, *C. elegans* encounters diverse microbial communities composed of phylogenetically related taxa with overlapping metabolic outputs. If attraction reflects recognition of a species-specific volatile signature, responses should not generalize to other bacteria. Alternatively, if animals detect conserved microbial metabolites, closely related species should elicit similar attraction. Distinguishing between these possibilities is essential for determining whether the olfactory system responds to bacterial identity or to broader ecological signals.

To determine whether attraction to *S. enterica* reflects a species-specific response or a broader bias toward related bacteria, we tested olfactory preferences for representative isolates from the natural *C. elegans* microbiome, CeMbio (26). In two-choice assays, wild-type animals were presented with paired *E. coli* lawns, one exposed to odor from a microbiome isolate.

Wild-type animals displayed strong attraction to odors from BIGb0393, CEN2ent1, and JUb66 (Figure 3A), three isolates belonging to the Enterobacteriaceae family. In contrast, odors from MYb11 and BIGb0172 elicited little or no preference, and MYb71 induced weak aversion. These results demonstrate that *C. elegans* discriminates among microbial odors.

**Figure 3.**
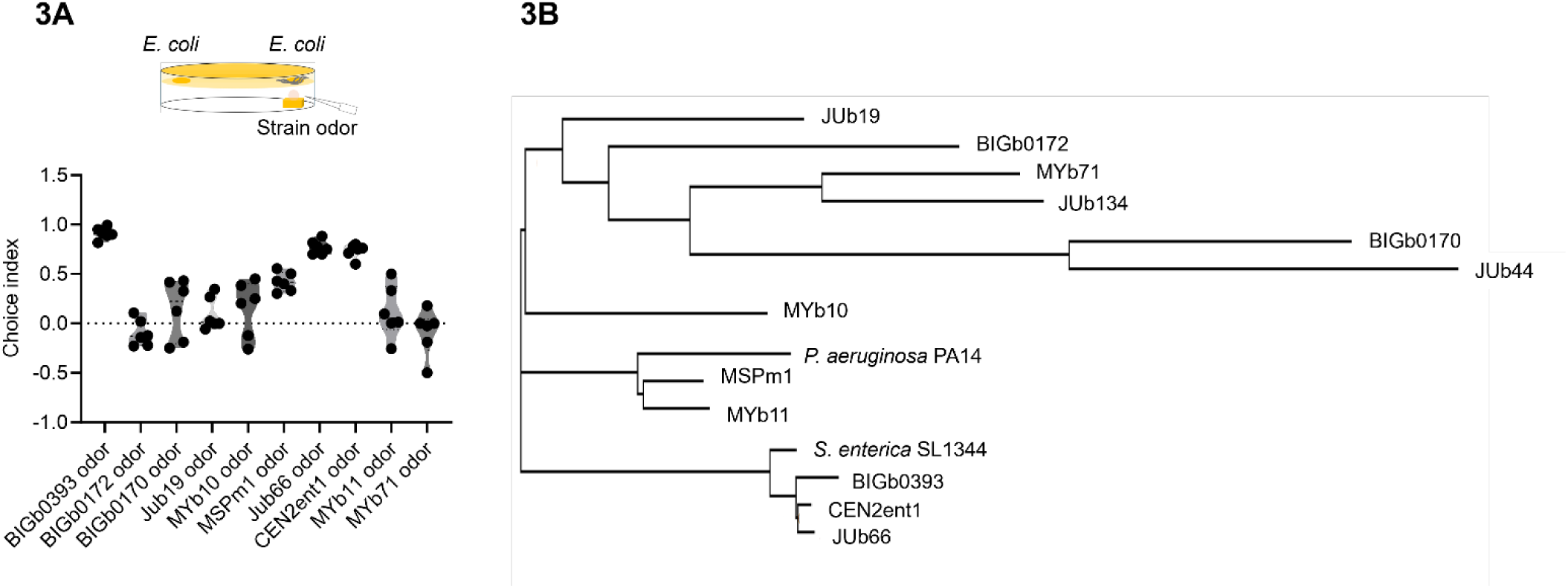
(A) Two-choice behavioral assay was used to assess the effect of microbiome-derived odors on *C. elegans* preference. Animals were placed equidistant from two *E. coli* lawns—one exposed to the odor of a specific microbiome strain (e.g., BIGb0393, BIGb0172, BIGb0170, Jub19, MYb10, MSPm1, Jub66, CEN2ent1, MYb11, MYb71). Animal preference was assayed in a two-choice behavior assay, with the Choice Index (CI). (B) Phylogenetic tree based on 16S ribosomal RNA gene sequences showing the evolutionary relationships among bacterial isolates used in odor exposure experiments with *C. elegans*.

To determine whether behavioral similarity correlated with phylogenetic relationships among the tested isolates, we performed 16S rRNA phylogenetic analysis. Phylogenetic reconstruction revealed tight clustering of *S. enterica*, JUb66, CEN2ent1, and BIGb0393 within Enterobacteriaceae (Figure 3B). Notably, all members of this class elicited attraction, whereas non-Enterobacteriaceae isolates did not. Thus, behavioral output paralleled phylogenetic clustering, suggesting that conserved metabolic repertoires within Enterobacteriaceae generate volatile profiles recognized as favorable.

The variation in attraction among closely related Enterobacteriaceae further indicates that *C. elegans* responds to differences in volatile blends rather than taxonomic identity alone. Together, these findings demonstrate that attraction to *S. enterica* reflects a broader sensory bias toward conserved Enterobacteriaceae metabolic signatures, linking bacterial phylogeny to host behavioral preference.

### Isoamyl Alcohol Overrides Pathogen-Induced Avoidance via AWC Neurons

The previous results established that *S. enterica* odor carries innate odor preference (Figure 1A), that this preference is experience-dependent (Figure 1B–C), and that attraction is conserved across related Enterobacteriaceae (Figure 3A–B). Together, these findings suggest that *C. elegans* responds to conserved metabolic outputs rather than species-specific identity. In fluctuating microbial environments, animals must balance nutrient acquisition against infection risk, implying that behavioral output reflects weighted integration of competing sensory cues. A strong, attractive odor could therefore override avoidance behavior if it reliably signals a favorable food source. We therefore asked whether a defined, conserved volatile metabolite is sufficient to reweight pathogen-induced avoidance.

Enterobacteriaceae produce the leucine-derived volatile isoamyl alcohol (IAA), a well-characterized attractant detected by AWC olfactory neurons (4, 27, 28). Because AWC activation promotes approach behavior, IAA allows us to test whether a single metabolite can dominate circuit output and counteract aversive memory. As expected, wild-type animals exhibited significantly greater occupancy on the *E. coli* lawn paired with IAA odor than with the control odor (Figure 4A). To test whether IAA alone can override learned aversion, Gravid adult wild-type (N2) animals were conditioned overnight on *P. aeruginosa* to induce avoidance and then transferred to a two-choice assay between *E. coli* and *P. aeruginosa* lawns with or without IAA odor. Despite prior pathogen exposure, *P. aeruginosa*-conditioned animals exhibited strong attraction to the IAA-paired lawn (Figure 4B), indicating that a single volatile metabolite is sufficient to reverse pathogen-induced avoidance.

**Figure 4.**
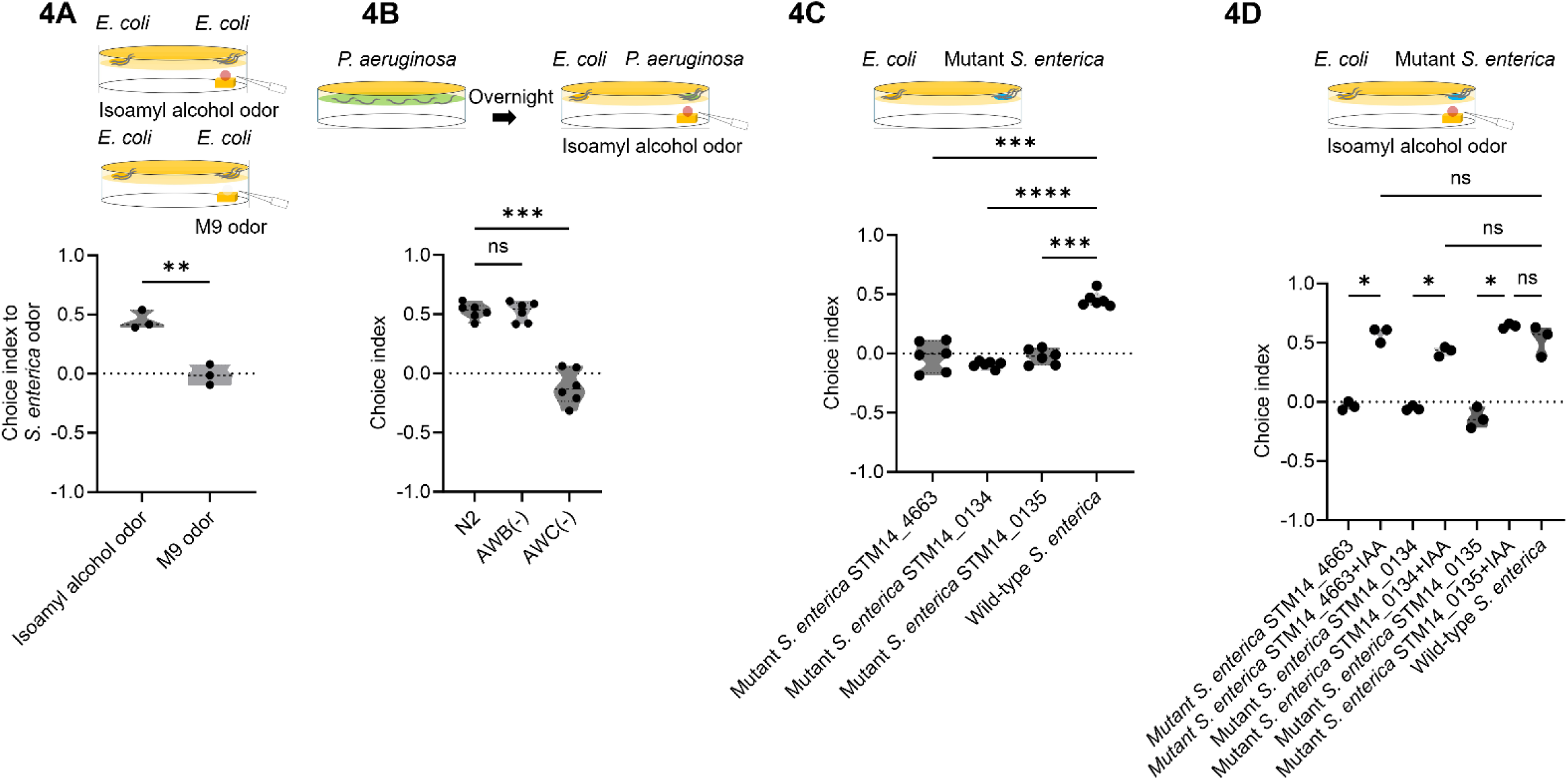
(A) Gravid adult *C. elegans* were given a choice between an *E. coli* lawn paired with Isoamyl alcohol odor and a *E. coli* lawn without odor. *t*-test was used and P values for CI of Isoamyl alcohol odor were calculated with respect to the M9 odor control. ****P < 0.0001, ***P < 0.001, **P < 0.01, and *P < 0.05. ns, non-significant. (B) Gravid adult *C. elegans* (wild-type N2, AWB (-), or AWC (-) strains) were preconditioned by feeding on *P. aeruginosa*. The following day, animals were subjected to a two-choice assay between *E. coli* and *P. aeruginosa* lawns supplemented with Isoamyl alcohol odor. Choice Index (CI) was calculated as: (number of animals on Isoamyl alcohol side – number on control side) / (number of animals on Isoamyl alcohol side + number on control side). CI > 0 indicates preference for the odor. One-way ANOVA with Tukey’s multiple comparisons test was used and P values for neuron–ablated animals were calculated with respect to the N_2_. ****P < 0.0001, ***P < 0.001, **P < 0.01, and *P < 0.05. ns, non-significant. (C) *C. elegans* were subjected to a two-choice assay between wild-type and mutant *S. enterica* strains (STM14_4663, STM14_0134, and STM14_0135). Animal preference was quantified using a Choice Index (CI), calculated as (number on mutant – number on *E. coli* lawn) / (number on mutant + number on *E. coli* lawn). CI values greater than zero indicate preference for the mutant strain, while values less than zero indicate preference for the *E. coli* lawn. One-way ANOVA with Tukey’s multiple comparisons test was used and P values for mutant *S. enterica* were calculated with respect to the wild-type *S. enterica*. ****P < 0.0001, ***P < 0.001, **P < 0.01, and *P < 0.05. ns, non-significant. (D) Gravid adult *C. elegans* were subjected to a two-choice assay comparing *S. enterica* mutant strains defective in the leucine biosynthesis pathway, which is responsible for Isoamyl alcohol (IAA) production. The tested strains were STM14_4663, STM14_0134, and STM14_0135. Bacterial lawns were supplemented with or without exogenous Isoamyl alcohol odor. Animal preference was quantified using a Choice Index (CI), calculated as (number on mutant strain lawn − number on OP50 lawn) / (number on mutant strain lawn + number on OP50 lawn). CI > 0 indicates preference for the test strain; CI < 0 indicates preference for the comparison strain. Error bars denote SD from six independent experiments. *t*-test was used and P values were calculated for each mutant *S. enterica* with respect to *S. enterica* supplemented with isoamyl alcohol odor. ****P < 0.0001, ***P < 0.001, **P < 0.01, and *P < 0.05. ns, non-significant.

To define the neuronal requirement for this override, we repeated the assay in sensory neuron–ablated strains. AWC ablation abolished attraction to the IAA-paired lawn, reducing preference to baseline, whereas AWB ablation had no effect (Figure 4B). Thus, AWC neurons are required for IAA-mediated suppression of avoidance. Together, these results demonstrate that IAA, acting through AWC neurons, functions as a positive signal capable of reweighting approach–avoidance decisions. Behavioral responses are not fixed by prior pathogen experience but instead reflect integration of competing sensory cues.

### Leucine-dependent isoamyl alcohol production is required for *S. enterica* attraction

The previous results identified isoamyl alcohol (IAA) as a candidate volatile cue contributing to attraction to *S. enterica*. To investigate whether bacterial leucine biosynthesis contributes to this behavioral preference, we examined *S. enterica* strains carrying mutations in *leuA* (STM14_0134), *leuL* (STM14_0135) and *atpA* (STM14_4663) mutant strains.

In two-choice assays between *E. coli* and leucine-pathway mutant *S. enterica* strains, *C. elegans* showed no preference between the *E. coli* and *S. enterica* mutant lawns (Figure 4C). Mutants lacking *leuA* (STM14_0134), *leuL* (STM14_0135), or *atpA* (STM14_4663) exhibited choice indices near zero, indicating loss of the attraction observed with wild-type *S. enterica*. The ATP synthase mutant *atpA* (STM14_4663), which exhibits a broad bioenergetic defect, was included as a general metabolic control. Together, these results demonstrate that leucine-dependent volatile production is required for the normal olfactory appeal of *S. enterica*.

To determine whether IAA is sufficient to restore attraction, we supplemented mutant lawns with exogenous IAA odor. IAA addition significantly rescued chemotaxis (Figure 4D). Thus, exogenously supplied IAA is sufficient to restore attraction in a mutant that has lost it. By producing IAA, *S. enterica* engages conserved olfactory pathways in *C. elegans*, specifically AWC-mediated attraction. In decomposing substrates, IAA levels likely correlate with bacterial growth and nutrient availability, making it a reliable ecological signal. The use of such conserved metabolic cues may allow pathogenic Enterobacteriaceae to exploit pre-existing host sensory biases, enhancing host contact despite potential fitness costs.

### Hydrogen sulfide contributes to *S. enterica* attraction

In natural environments, animals encounter complex odor blends rather than single compounds, and behavioral output can reflect integration of multiple bacterial volatiles (4, 5). If *C. elegans* relies on conserved indicators of microbial metabolic activity, other volatiles besides IAA should contribute to *S. enterica* attraction.

Hydrogen sulfide (H₂S) is a strong candidate to influence attraction, as it is produced through dissimilatory sulfur metabolism and cysteine degradation pathways in many Enterobacteriaceae, including *S. enterica*, and accumulates in decomposing substrates where bacterial density and metabolic activity are high (13, 14). Because H₂S is a product of bacterial sulfur metabolism, we hypothesized that it could serve as an additional microbial cue influencing C. elegans attraction.

We first tested whether H₂S is sufficient to promote attraction. In two-choice assays, *C. elegans* preferentially accumulated on an *E. coli* lawn paired with H₂S odor (Figure 5A), demonstrating that attraction was H₂S-dependent.

**Figure 5.**
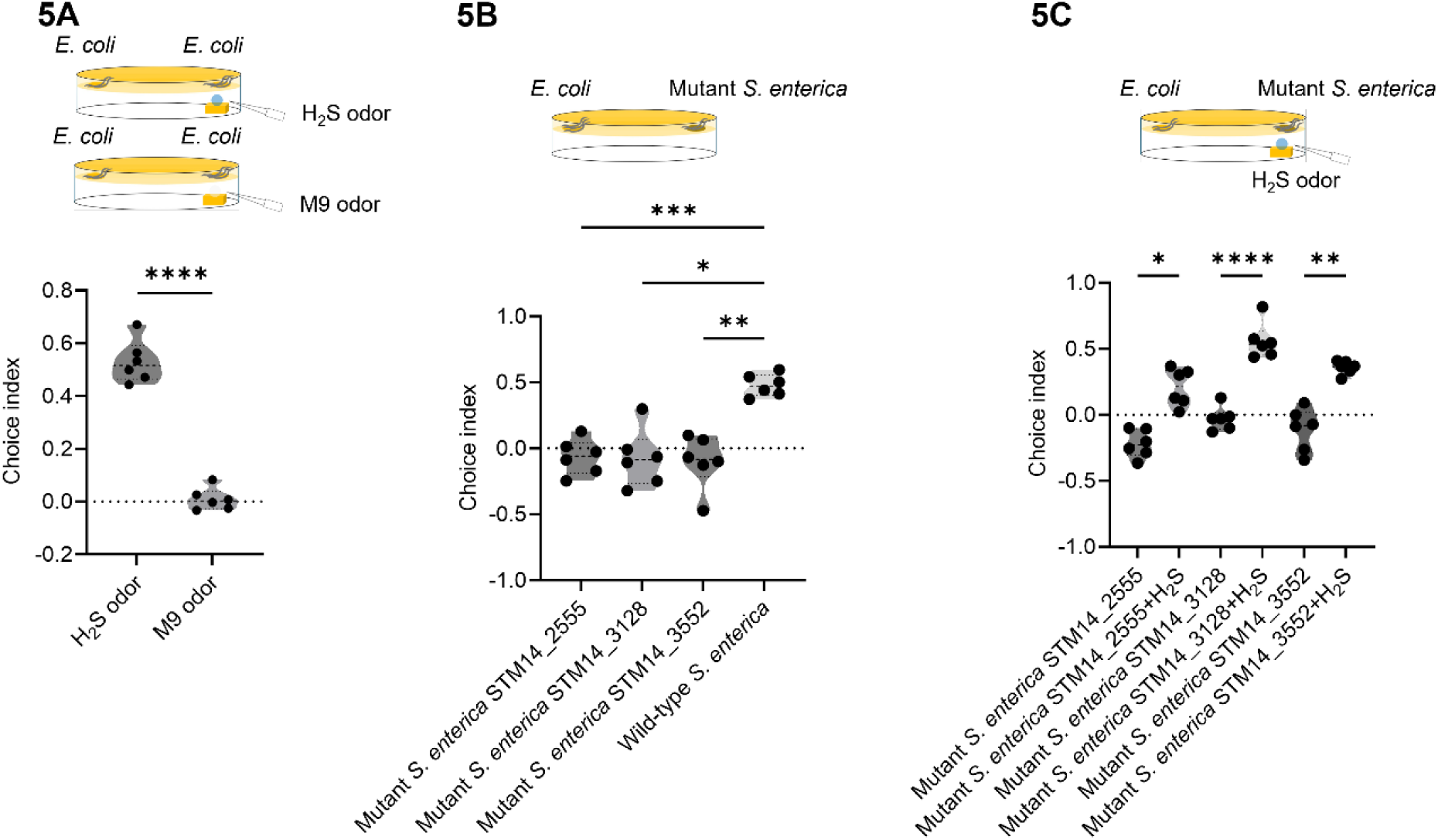
(A) Gravid adult *C. elegans* were given a choice between an *E. coli* lawn paired with hydrogen sulfide odor and an *E. coli* lawn without paired odor. *t*-test was used and P values for H_2_S odor were calculated with respect to the M9 buffer control. ****P < 0.0001, ***P < 0.001, **P < 0.01, and *P < 0.05. ns, non-significant. (B) *C. elegans* were subjected to a two-choice assay between wild-type and mutant *S. enterica* strains (STM14_2555, STM14_3128, and STM14_3552). Animal preference was quantified using a Choice Index (CI), calculated as (number on mutant – number on *E. coli* lawn) / (number on mutant + number on *E. coli* lawn). CI values greater than zero indicate preference for the mutant strain, while values less than zero indicate preference for the *E. coli* lawn. *t*-test was used and P values were calculated with mutant *S. enterica* respect to the wild-type *S. enterica*. ****P < 0.0001, ***P < 0.001, **P < 0.01, and *P < 0.05. ns, non-significant. (C) The Choice Index (CI) for *C. elegans* preference towards wild-type *S. enterica* (WT), STM14_2555 mutant, STM14_3128 mutant, and STM14_3552 mutant strains. Mutant bacterial lawns were supplemented with or without exogenous H₂S. CI values were calculated as (number on mutant lawn – number on *E. coli* lawn) / (number on mutant lawn + number on *E. coli* lawn), where a positive CI indicates attraction and a negative CI indicates aversion. *t*-test was used and P values were calculated with mutant *S. enterica* respect to the mutant *S. enterica* supplemented with H_2_S odor. ****P < 0.0001, ***P < 0.001, **P < 0.01, and *P < 0.05. ns, non-significant.

To determine whether bacterial H₂S production is required for *S. enterica* attraction, we examined mutants defective in key sulfur metabolism genes *PhsA* (STM14_2555), *asrC* (STM14_3128), and *cysI* (STM14_3552). Each of these mutants exhibited significantly reduced attraction relative to wild-type *S. enterica* (Figure 5B). Supplementation with exogenous H₂S odor restored attraction to mutant strains (Figure 5C), confirming that loss of this volatile accounted for the behavioral deficit. Notably, the sulfur-pathway mutants still produce IAA, and the leucine-pathway mutants still produce H₂S, yet neither are attractive (Figures 4C and 5B). Thus, although exogenous IAA or H₂S is individually sufficient to drive attraction, neither volatile is sufficient on its own at the levels present in the bacterium, indicating that at physiological concentrations, IAA and H₂S act together as a two-component signal. Together, they define a conserved metabolic signature characteristic of Enterobacteriaceae. These findings indicate that *C. elegans* evaluates microbial communities through core metabolic outputs, linking bacterial physiology to host foraging behavior.

## DISCUSSION

Volatile metabolites signal to an animal that bacteria are present and growing, but they do not indicate whether those bacteria are beneficial or a threat. Here we show that *C. elegans* is attracted to *S. enterica* and closely related Enterobacteriaceae through volatile metabolites, and that this behavior is mediated by a chemosensory circuit involving ASE, ASI, and AWC neurons. This response is not fixed but can be reassigned when *S. enterica* odor is encountered during exposure to pathogenic *P. aeruginosa*, indicating that olfactory-driven attraction is subject to experience-dependent modification. We further identify isoamyl alcohol (IAA) and hydrogen sulfide (H₂S) as key bacterial metabolites that drive *S. enterica* attraction, linking conserved aspects of bacterial metabolism to host behavior.

The attraction of *C. elegans* to a pathogenic bacterium contrasts with its well-established ability to avoid harmful microbes. Avoidance of *P. aeruginosa* emerges after exposure and infection and depends on physiological changes associated with intestinal colonization (16, 21). In contrast, volatile cues provide information at a distance before ingestion. These observations indicate that attraction and avoidance reflect distinct modes of sensory evaluation. Attraction to *S. enterica* is therefore consistent with the detection of cues that normally signal the presence of metabolically active bacteria, even when those cues originate from pathogens.

Attraction was not restricted to *S. enterica* but extended to multiple Enterobacteriaceae isolates, indicating that *C. elegans* responds to conserved features shared across related bacteria. Behavioral responses paralleled phylogenetic relationships, consistent with the detection of common metabolic outputs. IAA and H₂S, identified here as key drivers of attraction, are products of central metabolic pathways and are enriched in bacteria that proliferate in decomposing substrates. Together, these findings support a model in which *C. elegans* relies on conserved metabolic cues rather than species-specific signals to guide foraging decisions.

The behavioral response to these cues can be modified by experience. When *S. enterica* odor is paired with exposure to *P. aeruginosa*, attraction is reduced and, at early time points, reversed into avoidance, indicating that the response to a given odor can be reassigned based on prior exposure. This is consistent with previous work showing that *C. elegans* learns to avoid odors associated with pathogenic bacteria following infection ({29). However, in this case reassignment occurs at the level of volatile cues, before direct contact with the pathogen. This indicates that olfactory representations are not fixed but are shaped by the association between sensory input and internal state.

The requirement for ASE, ASI, and AWC neurons indicates that this reassignment is mediated by a distributed chemosensory network. AWC neurons detect volatile attractants, whereas ASE and ASI have been implicated in integrating environmental cues with physiological state and pathogen responses. These observations suggest that experience-dependent changes in odor preference arise from circuit-level integration rather than from modulation of a single sensory neuron class. Such integration is consistent with the broader organization of the *C. elegans* chemosensory system, in which multiple sensory modalities converge to regulate behavior.

A notable feature of our findings is that a single metabolite can override learned avoidance. IAA was sufficient to reverse aversion induced by *P. aeruginosa*, indicating that strong attractive signals can dominate behavioral output. This finding is consistent with a framework in which animals integrate competing sensory inputs, with behavioral responses reflecting the relative strength and reliability of available cues. In complex and dynamic environments, such flexibility would allow animals to balance the acquisition of nutrients with the risk of pathogen exposure.

Our results also raise the possibility that bacterial pathogens exploit conserved sensory biases in their hosts. By producing metabolites that are commonly associated with nutrient-rich conditions, pathogens such as *S. enterica* may promote host approach and increase the likelihood of ingestion. Similar dynamics have been described for *P. aeruginosa*, which can initially attract *C. elegans* but subsequently induces learned avoidance following infection (21). More broadly, bacterial metabolites can act as attractive or aversive foraging cues in this nematode (32, 33). The use of conserved metabolic signals may therefore represent a general mechanism through which microbes influence host behavior.

In summary, we show that two conserved products of bacterial metabolism drive attraction to a pathogen, that this attraction is read out through a defined chemosensory circuit, and that experience can reassign the meaning of the odor. An attractive smell is therefore not a fixed property of a bacterium but a judgment the nervous system makes and can revise.

## MATERIALS AND METHODS

### Bacterial strains

The bacterial strains used in this study included *Escherichia coli* OP50, *Pseudomonas aeruginosa* PA14, *Salmonella enterica* serovar Typhimurium 14028, and the *S. enterica* mutant strains STM14-4663, STM14-0134, STM14-0135, STM14-2555, STM14-3128, and STM14-3552 (33). CeMbio strains were obtained from the *Caenorhabditis elegans* Genetics Center (CGC) and included *Sphingobacterium multivorum* BIGb0170, *Acinetobacter guillouiae* MYb10, *Lelliottia amnigena* JUb66, *Stenotrophomonas indicatrix* JUb19, *Pseudomonas lurida* MYb11, *Pseudomonas berkeleyensis* MSPm1, *Comamonas piscis* BIGb0172, *Pantoea nemavictus* BIGb0393, *Ochrobactrum vermis* MYb71, and *Enterobacter hormaechei* CEN2ent1.

### C. elegans strains

The following *C. elegans* strains used in this study were obtained from the CGC: N2 (Wild-Type strain), PR680 che-1(p680), PY7505 oyIs84 [gpa-4p::TU#813 + gcy-27p::TU#814 + gcy-27p::GFP + unc-122p::DsRed], JN1715 peIs1715 [str-1p::mCasp-1 + unc-122p::GFP], PY7502 oyIs85 [ceh36p::TU#813 + ceh-36p::TU#814 + srtx-1p::GFP + unc-122p:DsRed].

### Aversive training

Training plates of 6-cm-diameter containing full-lawns of *P. aeruginosa* on SK agar (modified NGM agar plates [0.35% instead of 0.25% peptone]). Gravid adult hermaphroditic animals grown on *E. coli* OP50 were washed with M9 and transferred to the training plates overnight They were then transferred to the appropriate assay plates.

### Odor exposure

Bacterial cultures were grown by inoculating individual bacterial colonies into 2 mL of LB broth and growing them for 12hr on a shaker at 37°C. Bacterial odor was delivered by placing a 30 µL aliquot of culture on an agar plug on the lid of the assay plate. Plates were then inverted to prevent contamination of the agar surface without direct contact with the animals while permitting diffusion of airborne cues. Exogenous volatile exposure for Isoamyl alcohol (IAA) and hydrogen sulfide (H₂S) were freshly prepared in M9 buffer prior to each experiment. Isoamyl alcohol was diluted 1:1000, and hydrogen sulfide was diluted 1:2000. 5 μL of the diluted solution was applied on an agar plug on the lid of the assay plate. Plates were then inverted to prevent contamination of the agar surface without direct contact with the animals while permitting diffusion of airborne cues.

### Two-choice preference assay

Bacterial cultures were grown by inoculating individual bacterial colonies into 2 mL of LB broth and growing them for 12hr on a shaker at 37°C. Then, 20 µL of each was plated on opposite sides of a 6-cm-diameter SK plate and incubated overnight at 37°C. The plates were cooled to room temperature for at least 30 min before seeding with animals. Gravid adult hermaphroditic animals were transferred to the center of plates equidistant from both the lawns. The numbers of animals on both lawns were counted at the specified times for each experiment. Six 6-cm-diameter plates were used per trial in every experiment. The *S. enterica* choice index (*S. enterica* CI) was calculated as follows:

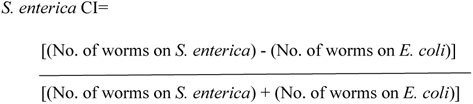

Choice indices to other bacteria were similarly calculated.

### Phylogenetic analyses

Phylogenetic analysis was performed using 16S rRNA gene sequences from 14 bacterial isolates. Sequences were aligned in MegAlign Pro (DNASTAR Lasergene 17.0.2.1) with the Clustal Omega algorithm, and global gap removal was applied before tree construction. Phylogenetic relationships were inferred using the BIONJ method with the Tamura–Nei (1993) distance metric. The resulting tree was displayed as an unrooted phylogeny.

### Statistical analysis

The statistical analysis was performed with Prism 10 (GraphPad). For violin plots, the center line depicts the median. For all violin plots, individual dots represent individual repeat. All experiments were performed in six biological repeats. Unpaired t-tests, one-way or two-way ANOVA with subsequent group comparisons were performed as indicated in the figure legends. In the figures, ns denotes not significant, and asterisks (*) denote statistical significance as follows: *P < 0.05; **P < 0.01; ***P < 0.001; ****P < 0.0001, as compared with the appropriate controls.

## ACKNOWLEDGMENTS

We thank Dr. Michael McClelland (University of California, Irvine) for generously providing the *Salmonella enterica* mutant strains used in this study. Most *Caenorhabditis elegans* strains used in this study were obtained from the *Caenorhabditis* Genetics Center (CGC), which is funded by the National Institutes of Health Office of Research Infrastructure Programs (P40 OD010440). This work was supported by the National Institutes of Health Grant GM070977.

## Notes

### Competing Interest Statement

The authors have declared no competing interest.

